# Probing voltage dependence interaction of cationic peptides with bacterial porins at a single-molecule level

**DOI:** 10.64898/2026.04.08.717161

**Authors:** Sonal Prasad

## Abstract

This study investigates the interaction between the cationic antimicrobial peptide protamine and bacterial porin OmpF (*E. coli*) at the single-molecule level. Using high-resolution conductance measurements in planar lipid bilayers, strong voltage- and concentration-dependent ion current blockages with OmpF, indicating significant protamine binding were observed. Further analysis revealed that peptide length influences binding kinetics, with longer peptides showing reduced affinity and slower exchange rates. These findings demonstrate that OmpF is a tractable model for studying cationic peptide–channel interactions and translocation mechanisms relevant to antimicrobial action.

## Introduction

The rise of antimicrobial resistance has intensified the search for novel therapeutic strategies, among which antimicrobial peptides (AMPs) have emerged as promising candidates (1-3). These peptides, particularly cationic antimicrobial peptides (CAPs), offer broad-spectrum activity and a reduced likelihood of resistance development. CAPs typically consist of 12–50 amino acids and, despite their sequence and structural diversity, share common features: a net positive charge (+4 to +6), a high proportion of hydrophobic residues (50-70%), and the ability to adopt amphipathic α-helical conformations in lipid environments (4, 5).

Protamine (Ptm) is a particularly intriguing CAP, composed of 21 arginine residues, making it the most highly cationic naturally occurring peptide reported to date. With a molecular weight of ∼4.2 kDa and an isoelectric point of ∼11–13, Ptm lacks a defined secondary structure due to the uniform distribution of positive charges along its backbone (6).

CAPs are generally believed to exert their antimicrobial effects by disrupting bacterial membranes, leading to ion leakage, membrane depolarization, and cell death. Several models have been proposed to describe this process, including the barrel-stave, toroidal pore, and carpet models (5, 7). AMP selectivity for prokaryotic cells is largely governed by non-receptor-mediated mechanisms, driven by recognition of the membrane’s physicochemical properties (8-10). However, microbial inactivation has also been observed at sub-lytic concentrations, suggesting that AMPs may act through intracellular mechanisms, such as inhibition of cell wall synthesis, transcription, translation, nucleic acid synthesis, enzymatic activity, or cytoplasmic aggregation (8, 11-15).

Recent findings indicate that Ptm can internalize into certain Gram-negative bacteria without disrupting their membranes. Moreover, Ptm does not perturb reconstituted bacterial membranes composed of native phospholipid mixtures, implying a non-lytic, protein-mediated translocation mechanism. One plausible pathway involves porin channels, which are abundant, non-specific membrane proteins in Gram-negative bacteria (16).

Among these, OmpF in Escherichia coli is a well-characterized porin, present in over 10^5^ copies per cell. It forms a trimeric complex of β-barrel structures, each composed of 16 antiparallel β-strands. A key structural feature is loop L3, which folds into the channel to form a constriction zone. This region is defined by a transversal electric field, generated by negatively charged residues (D113, E117) on the L3 side and positively charged residues (R42, R82, R132) on the opposite side (17-19).

Given the importance of charge interactions in peptide translocation, we hypothesized that cationic peptides such as Ptm exploit the electrostatic landscape of OmpF porins to facilitate their uptake (20). To test this, single porin channels were reconstituted into artificial planar lipid bilayers and peptide interactions using time-resolved ion current blockage measurements were monitored. External electric field modulated the dynamics of peptide entry and exit through the pore.

In this study, Ptm translocation across the outer membranes of *E. coli* and *S. typhimurium* via porin-mediated pathways were studied, despite its relatively large size compared to known porin-translocating antibiotics. The translocation of smaller cationic α-helical peptides through OmpF using a chip-based automated patch clamp technique were examined, enabling high-resolution analysis of rapid translocation events. How peptide charge and length influence the free energy barrier for transport across the OmpF pore were explored.

## Materials and methods

### Peptides

The long highly cationic peptides used in this study were protamine sulfate (ARRRRSSSRPIRRRRPRRRTTRRRRAGRRRR, Y1 from Clupeine P4505, Mw: 4110.51 Da), (Poly-L-lysine hydrobromide 25988-63-0, Mw: 500-2000 Da) and poly-arginine (Poly-L-arginine hydrochloride 26982-20-7, Mw: 5000-15000 Da), protamine C-terminal end (RAGRRRR). These synthetic peptides were ordered from Sigma-Aldrich AG, Germany. The short cationic peptides used in this study were penta-lysine (H-Lys-Lys-Lys-Lys-Lys-OH acetate salt, 19431-21-1, Mw: 658.88 Da), tetra-arginine (H– Arg–Arg–Arg–Arg–OH acetate salt 26791-46-8, Mw: 642.77 Da) and tri-arginine (H–Arg–Arg–Arg–OH acetate salt 19431-21-1, Mw: 486.54 Da). These synthetic peptides were ordered from Bachem AG, Switzerland.

### Planar lipid bilayer (BLM) preparation and single channel electrophysiological recordings

Solvent-free lipid bilayers were prepared using a modified version of the Montal–Mueller technique (23). A small aperture (∼60–80 µm in diameter) was created in a 25 µm thick Teflon film using a discharge spark (10–50 kV at 500 kHz) generated by a BD-10 Vacuum Tester. The Teflon film was then sealed with silicone grease between two glass half-cells (2.5 ml each), electrically isolating the aqueous compartments.

Prior to bilayer formation, the aperture was pre-treated with 1 µl of 1% n-hexadecane in n-hexane and allowed to dry for 10 minutes. Both chambers were subsequently filled with buffer solution (1 M KCl, 20 mM MES, pH 6.0). A lipid bilayer was formed by spreading 1 µl of 1,2-diphytanoyl-sn-glycero-3-phosphocholine (Avanti Polar Lipids, Alabaster, AL) from a 5 mg/mL solution in n-pentane onto the buffer surface. Bilayer formation occurred within 10 minutes as the solvent evaporated.

Ag/AgCl electrodes were used to record ionic currents, with the cis-side electrode grounded and the trans-side electrode connected to the headstage of an Axopatch 200B amplifier (Axon Instruments, Foster City, CA). Purified, detergent-solubilized porins (1 µl) were added to the cis chamber and inserted into the bilayer by applying a voltage of 150–200 mV. Successful insertion of single porins was confirmed by discrete increases in current.

For OmpF porins, the protein was diluted from a 1.5 mg/ml stock solution in 1% Octyl-POE detergent (Alexis, Lauchringen, Switzerland). Single-channel insertion typically occurred within 15 minutes, depending on protein concentration (21). The output signal was filtered using a low-pass Bessel filter at 10 kHz and sampled at 50 kHz (22, 23).

Recordings involving smaller peptides, such as triarginine, were performed using automated planar patch-clamp systems with microstructured glass chips. These integrated devices required <10 µl of solution and enabled rapid fluid exchange (24). The resulting bilayers exhibited low capacitance and improved signal-to-noise ratios, allowing high-resolution single-channel recordings (See, Supplementary text). All experiments were conducted at room temperature. Data analysis was performed using Clampfit software (Axon Instruments, Inc.).

### Determination of binding affinities and kinetics constants

Artificial lipid bilayers were formed using the BLM technique, into which single OmpF channels were reconstituted. The interaction of peptides with the channel was studied using single-channel electrophysiology, allowing the calculation of kinetic rate constants. This approach enabled the determination of peptide binding kinetics, including association and dissociation rate constants, as well as partitioning behavior.

Single-channel recordings of the OmpF pore revealed that peptide partitioning kinetics were influenced by two main factors: Increased event frequency with higher transmembrane potentials, indicating enhanced peptide interaction. Reduced event frequency for longer peptides, suggested hindered translocation through the nanometer-scale pore.

In these measurements, key parameters included: Residence time (*τ*_*r*_): Duration of peptide-induced blockages. Open channel time (*τ*_*o*_): Time between blockage events. Blockage frequency (*v*_*m*_): Number of blockage events per second per monomer (25).

The partition coefficient (P) between the aqueous phase and the pore lumen was derived from the equilibrium association constant (*K*_*f*_), assuming low pore occupancy (26):

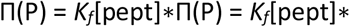

where the effective molar concentration of a single peptide inside the OmpF pore was:

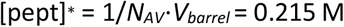

Here, *N*_*AV*_ is Avogadro’s number and *V*_*barrel*_ is the internal volume of the β-barrel (∼0.7 × 1.1 nm^2^).

To obtain chemical kinetic rates, equilibrium binding constants, and net substrate flux, peptide translocation was modeled as a chemical adsorption–desorption process between the substrate (S) and the pore (P). In cases of strong binding relative to diffusion, the interaction was treated as a reversible reaction: S + P ⇌ SP ⇌ P + S

This framework allowed quantification of binding dynamics and facilitated mechanistic insights into peptide transport through membrane channels.

## Results

We employed high-resolution ion conductance measurements to quantify peptide–channel interactions using a single trimeric porin reconstituted into a planar lipid bilayer.

### Interaction of Protamine with OmpF channel

Addition of the large cationic peptide 10 µM protamine (Ptm) to the cis (extracellular) side of the membrane required trans-negative potentials (−100 mV) to induce interaction with the OmpF channel. When Ptm was added to the trans side, voltage dependence was reversed: positive potentials increased event frequency, while negative potentials produced no current (Fig. 1Ai, ii). No binding events were observed when the electric field was reversed, positive potentials on the trans (intracellular) side, indicating voltage-dependent interaction in 1M KCl, 20 mM MES, pH 6.0 buffer (Fig. 1Aii). At higher transmembrane potential -100 mV, OmpF channel exhibited gating behavior, characterized by stepwise closure of the three monomers, in the presence of 10 µM of Ptm (Fig. 1Aiv) while at lower concentration of 5 µM Ptm no complete closure of the channel with mixed long and short blockage events was observed (Fig. 1Aiii, inset show mixed events). On applying negative potential the peptide interaction was prominent whereas on reverse potential no binding effect was noticed. Protamine at lower concentration exhibited weaker and shorter partial blockage events with the OmpD channel under the same conditions (Fig. 1Biii, inset show mostly short-lived events). At higher concentration Ptm displayed short and large events led to either transient blockage or complete channel closure, implying peptide internalization but not definitive translocation (Fig. 1Biv). No gating behaviour of both the channels were observed in the absence of peptide (Fig. 1Ai, Bi). Event frequency and residence time increased with both peptide concentration and applied voltage for OmpF and OmpD channels (Fig. S1A-C), further supporting voltage-dependent binding.

**Fig. 1.**
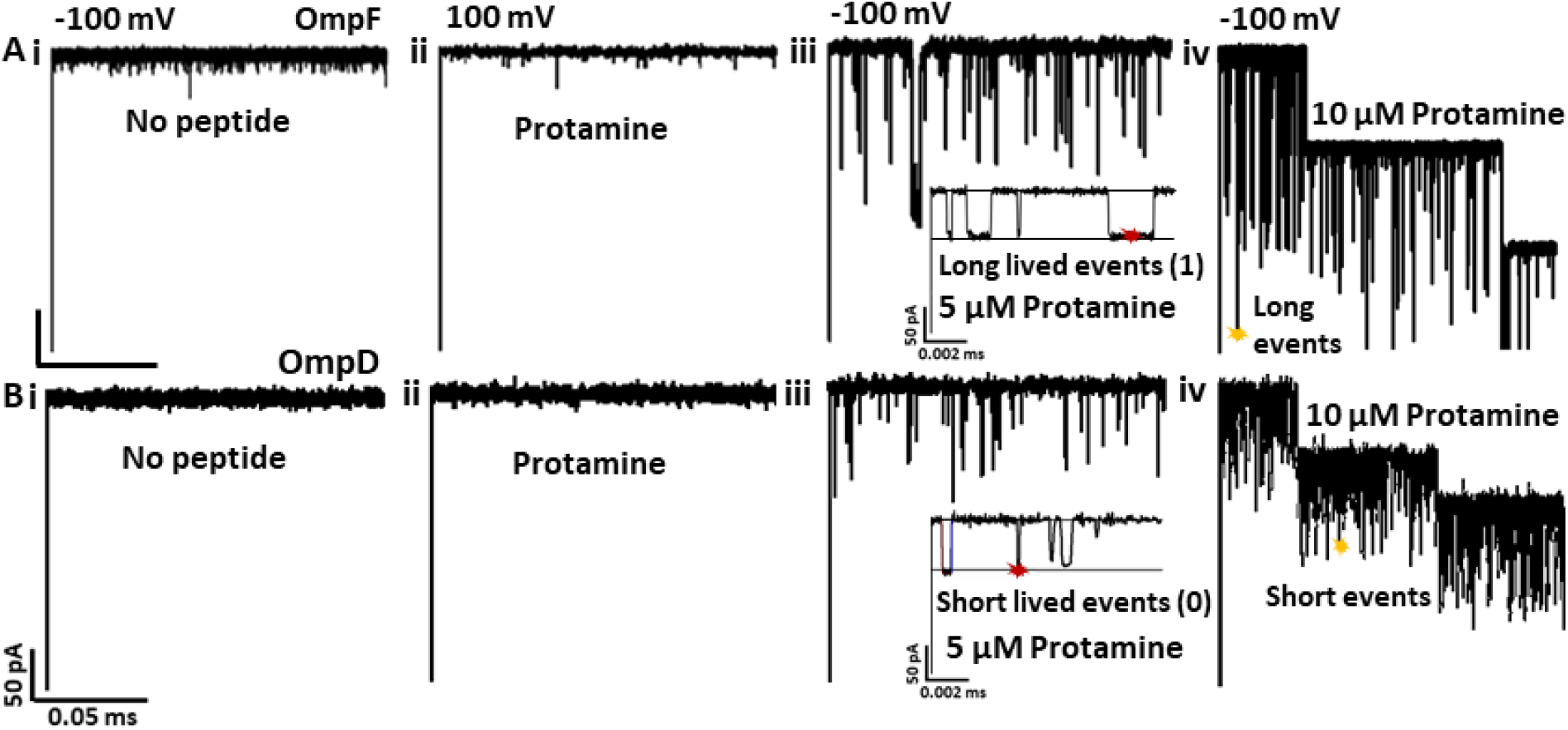
Typical ion current recordings through OmpF and OmpD porins with Protamine added to the cis side. (**A, B**) In the absence of peptide at -100 mV (**i**), in the presence of Protamine peptide showing no interaction at a reversal transmembrane potential of 100 mV (**ii**) through single trimeric OmpF and OmpD channels reconstituted into planar lipid membranes. (A, **iii**) 5 µM of Protamine on OmpF showing mixed long and short partial short blockage events at -100 mV. Inset showing long lived blocking events assigned (red star, 1). (A, **iv**) 3-step complete closure of OmpF pore displaying long events (yellow star) after applying 10 µM of Protamine at -100 mV. (B, **iii**) 5 µM of Protamine on OmpD showing short partial short blockage events at -100 mV. Inset show short lived blocking events assigned (red star, 0). (**B, iv**) 3-step closure of OmpD pore after applying 10 µM of Protamine with short events (yellow star) at -100 mV. Experimental conditions: T = 25°C, 1 M KCl, 20 mM MES, pH = 6.

To confirm the specificity of Protamine interaction with OmpF porin and understand the role of specific amino residues involved in peptide interaction, available mutants of the channels were used. Negatively charged residues in the L3 loop of the OmpF channel were examined through double mutant OmpF (D113N-E117Q) and single mutant OmpF (E117Q). Recordings with Ptm showed complete (Fig. S2A) and partial channel blockage (Fig. S2B). Recording with short cationic Tetra-arginine peptide on double mutant OmpF as a control showed no blockage events (Fig. S2C) suggesting that the two amino acid resides play a significant role for protamine interaction with the channel through binding with those residues.

Additional measurements using dynamic light scattering, UV spectroscopy and fluorescence spectroscopy by performing liposome swelling assay, Zeta potential size measurement and substrate selective supramolecular tandem assay interaction of protamine with OmpF channel was confirmed (Fig. S3 and S4). POPC liposomes reconstituted with OmpF porins showed change in the optical density (rate of swelling) and Osmolality after addition of Protamine (27)(Fig. S3A). Through Zeta potential measurements the size of the liposomes reconstituted with OmpF changed after addition of Protamine (Fig. S3B). In fluorescence spectroscopy measurements a complete displacement of the fluorescent LCG dye from the macrocyclic host CX4 cavity by the competitor protamine was determined through change in fluorescence intensity (28) (Fig. S4Ai). Liposomes reconstituted with OmpF with the integrated host-dye molecule (29) inside confirmed the passing of the protamine peptide through the OmpF porin and effectively displacing the dye from the host through change in the fluorescence intensity (Fig. S4Aii). Additional recordings in 0.1 mM KCl and 100 mM KCl buffer helped in better understanding of protamine interaction with the porins. In 0.1 M KCl, 20 mM MES, pH 6.0 buffer, the OmpF channel displayed a rapid closure on addition of protamine at -100 mV.

Interaction of longer cationic peptides such as poly-arginine and poly-lysine with OmpF channel was examined. Both peptides induced mixed long and short blockage events, with two-step channel closure from poly-arginine (Fig. 2A, B; insets show long and short-lived events). Two distinct types of blockade events, short-lived and long-lived were detected, indicating a strong interaction with the OmpF channel similar to Protamine. Additional measurement was conducted using the C-terminal fragment of protamine, comprising seven amino acid residues (RAGRRRR). The peptide was enzymatically digested with thermolysin and subsequently purified. Notably, the C-terminal fragment exhibited effects comparable with poly-lysine and other short cationic peptides indicating a monomer-specific interaction with the OmpF channel (Fig. 2C).

**Fig. 2.**
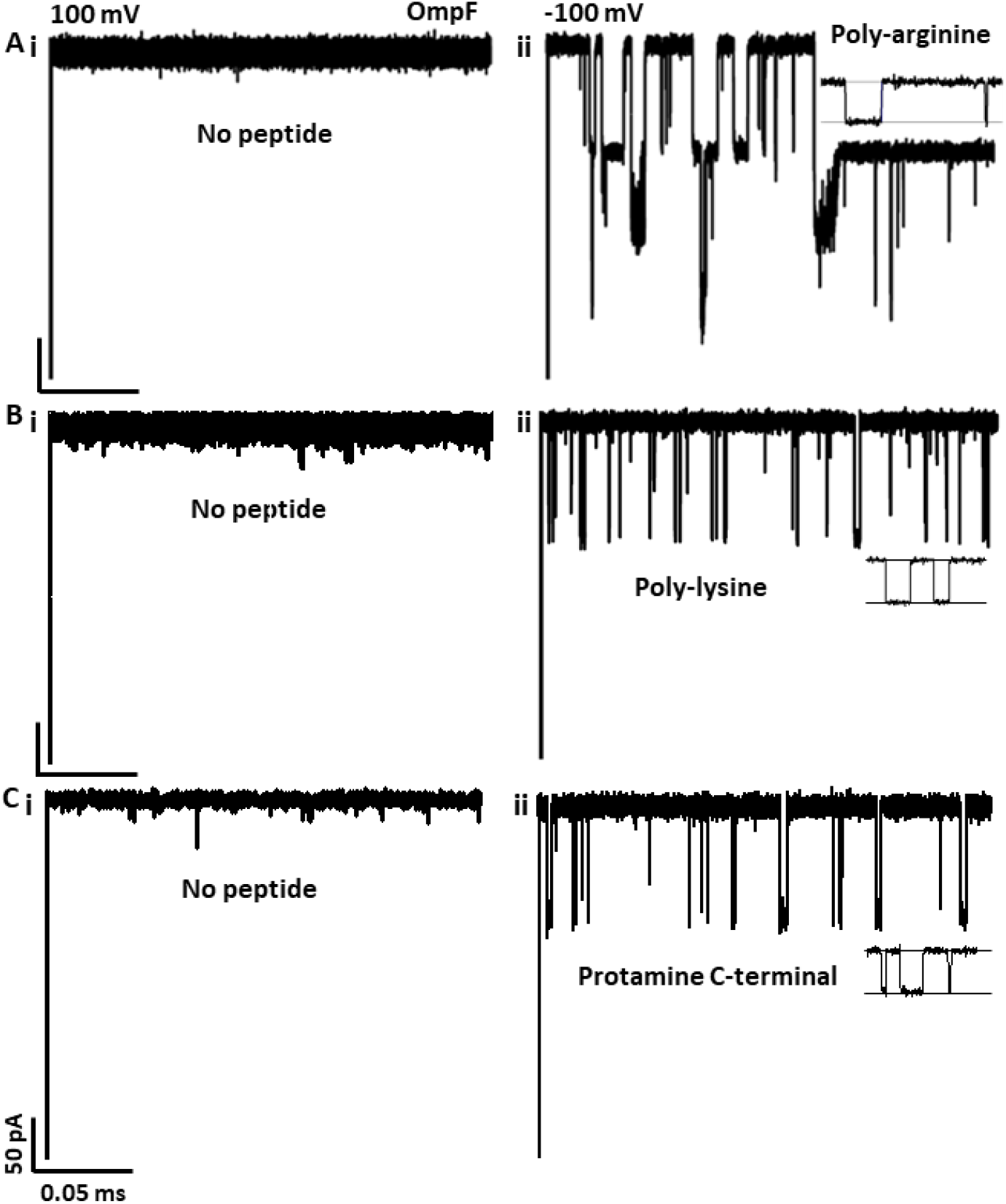
Typical ion current recordings through OmpF porin with large cationic peptides added to the cis side. (**A, B, C**) In the absence of peptides at a reversal transmembrane potential of 100 mV through single trimeric OmpF channels reconstituted into planar lipid membranes (**i**). (**A, ii**) 5 µM of Polyarginine showing effective gating of channel with two-step closure displaying mixed long and short blockage events. (**B, ii**) 5 µM of Polylysine displaying stronger interaction with mostly short blockage events. (**C, ii**) 5 µM of Protamine C-terminal end displaying mostly short blockage events. Insets show mixed long and short-lived blocking events. Current recording with peptides performed at -100 mV. Experimental conditions: T = 25°C, 1 M KCl, 20 mM MES, pH = 6.

The data demonstrates that protamine strongly interacts with the OmpF porin, leading to complete channel closure at -100 mV.

### Transient current blockades of the OmpF channel by cationic short peptides

Measurements were performed using short cationic peptides penta-lysine, tri-arginine, and tetra-arginine to investigate their potential translocation through the β-barrel OmpF pore via similar ion current fluctuation recordings. As previously noted, peptide-channel interactions were influenced by the side of peptide addition and the applied transmembrane potential.

When peptides were added to the cis side of the bilayer, negative potential significantly increased the frequency of blockade events, whereas positive potential did not induce any detectable blockages. The addition of peptides of varying lengths to the cis side produced reversible channel blockades at transmembrane potentials of -50 mV and -100 mV (Fig. 3). These blockades manifested as ion current fluctuations, indicating strong interactions with the OmpF channel, albeit with distinct binding kinetics. At 5 µM peptide concentration, monomer-specific blocking events were observed in single trimeric OmpF channels (Fig. 3A, B, Ciii). Penta-lysine and tetra-arginine induced complete monomer blockages, while tri-arginine caused partial closures. In fluorescence spectroscopy measurements an effective displacement of the fluorescent LCG dye from the macrocyclic host CX4 cavity by the competitor penta-lysine and tri-arginine was tested through change in fluorescence intensity (Fig. S4B-Ci). Liposomes reconstituted with OmpF with the integrated host-dye molecule inside showed a possibility of peptides displacing the dye from the host through change in the fluorescence intensity (Fig. S4B-Cii).

**Fig. 3.**
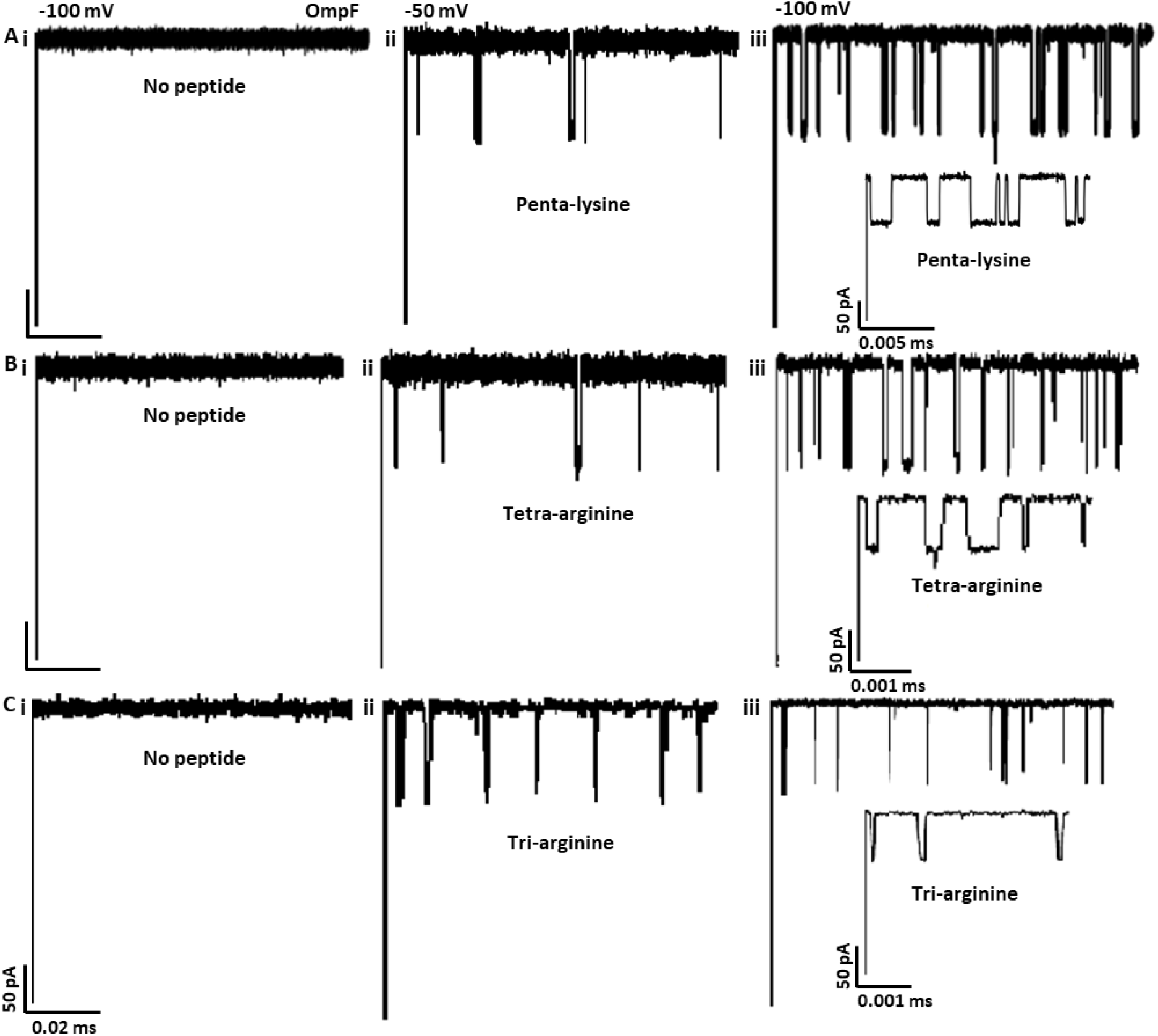
Typical ion current recordings through OmpF porin with small cationic peptides added to the cis side. (**A, B, C**) In the absence of peptides at a transmembrane potential of -100 mV through single trimeric OmpF channels reconstituted into planar lipid membranes (**i**). (A, B, C) 15 µM of Penta-lysine, 3 µM of Tetra-arginine, and 10 µM of Tri-arginine displaying short blockage events at -50 mV (**ii**). (A, B, C) All three long cationic peptides at -100 mV displayed short blockage events with mixed long and short -lived events (**iii**). Insets show single monomers short and long blocking events for all peptides. Experimental conditions: T = 25°C, 1 M KCl, 20 mM MES, pH = 6.

Dwell time analysis of the blockade events (fitted using a double exponential decay model) revealed the interaction strength hierarchy: penta-lysine > tetra-arginine > tri-arginine (Fig. 4). Both penta-lysine and tetra-arginine exhibited larger current amplitudes and longer dwell times compared to tri-arginine (Fig. 4A-Cii, iii). Protamine C-terminal end exhibited larger current amplitudes and longer dwell times (Fig. S2D). Notably, two distinct blockade peaks were observed: peak 0 (short-lived events) and peak 1 (long-lived events). The occurrence of peak 1 was dependent on voltage and peptide length, whereas peak 0 appeared consistently across all tested transmembrane potentials. Peak 0 events were characterized by low-amplitude, short-duration blockades, while peak 1 events displayed high-amplitude, long-duration blockades with higher frequency (Fig. 4, 5), suggesting partial peptide entry into the OmpF channel without traversing from cis to trans side of the pore. Semilogarithmic *τ*_*on*_ histograms were fitted with single exponential functions for all the three peptides, and event frequencies were normalized to the -50 mV transmembrane potential (Fig. 4). The amplitude of peak 1 blockades implied full partitioning of the peptide into the OmpF channel (28). The peaks represented transient current blockades induced by peptide addition to the cis side of the bilayer (Fig. 5). No blockade events were observed with 5 µM tetra-arginine on the double mutant OmpF (D113N-E117Q), indicating that amino acid residues D113 and E117 were critical for Ptm peptide binding (Fig. S2C). The data suggests that moderate length peptides interacted near the constriction zone and were more likely to traverse the OmpF pore compared to shorter peptides.

**Fig. 4.**
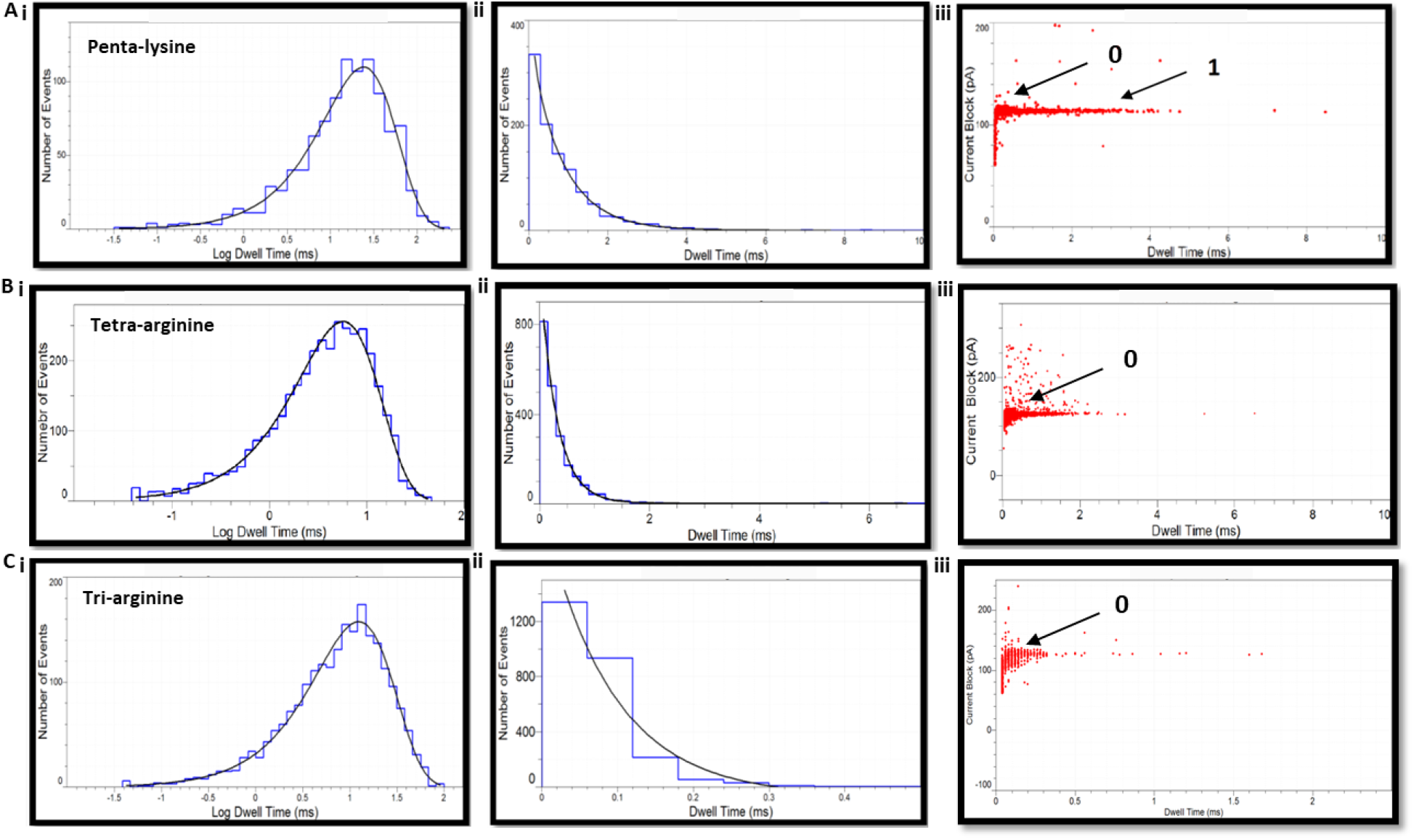
Semilogarithmic histogram and scatter plot of polypeptides interaction with OmpF pore. (**A, B, C**) Semilogarithmic histogram of inter-event intervals of Penta-lysine (*τ*_*on*_ = 29.87 ± 1.19ms), Tetra-arginine (*τ*_*on*_ =5.78 ± 0.11 ms) and Tri-arginine (*τ*_*on*_ = 12.41 ± 0.02 ms) fitted with Gaussian fit (**i**). (A, B, C) Dwell-time histogram of Penta-lysine (*τ*_*off*-0_ = 0.10 ± 0.20 ms, *τ*_*off*-1_ = 0.83 ± 0.04 ms), Tetra-arginine (*τ*_*off*-0_ = 0.28 ± 0.05 ms, *τ*_*off*-1_ = 0.57 ± 0.45 ms), Tri -arginine (*τ*_*off*-1_ = 0.09 ± 0.02 ms) fitted with single exponential fit (**ii**). (A, B, C) Scatter plot of current block amplitudes versus dwell time of peptides (**iii**) added to the cis side at an applied voltage of-100 mV.

**Fig. 5.**
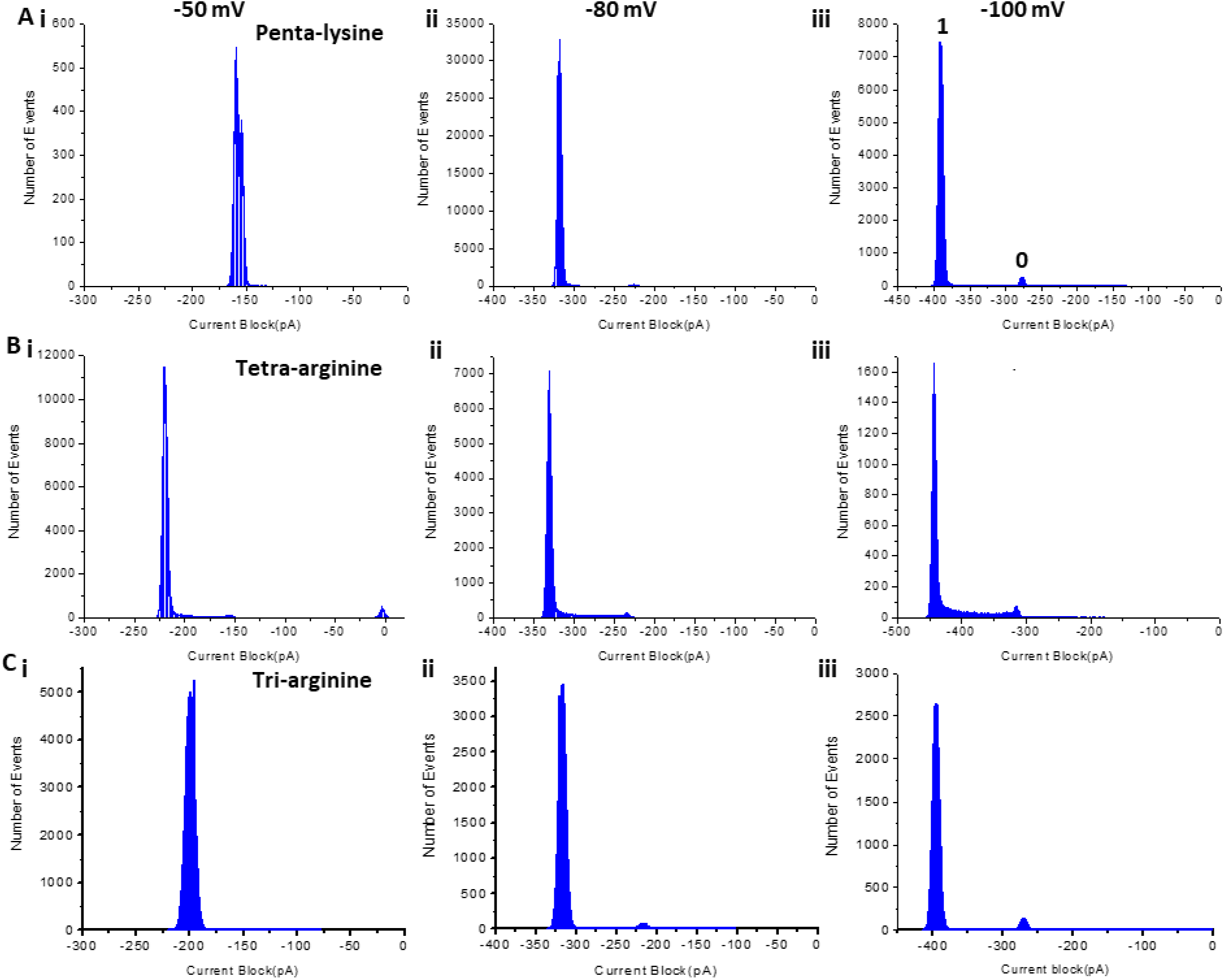
Typical amplitude histograms showing transient single-channel blockage of the OmpF pore voltage dependent of short cationic peptides. (**A**) 15 µM Penta-lysine, (**B**) 3 µM Tetra-arginine and (**C**) 10 µM Tri-arginine added to the cis side of single trimeric OmpF channels reconstituted into planar lipid membranes at a transmembrane potential of -50 mV (**i**), -80 mV (**ii**) and -100 mV (**iii**). At - 100 mV all the three short peptides showed major (1) and minor peaks (0). Experimental conditions: T = 25°C, 1 M KCl, 20 mM MES, pH = 6.

For tri-arginine, only short-lived blockade events were detected, with residence times around 100 µs. This highlights the limitations of classical planar lipid bilayer setups in capturing fast-binding events. To overcome this, a chip-based automated patch-clamp bilayer technique was employed. The miniaturized setup reduced background noise and electromagnetic interference, enabling low-noise recordings optimal for trimeric OmpF channels. Bilayers were formed via giant liposome adsorption, requiring minimal sample volumes (3-5 µl electrolyte drops on either side of the glass aperture; (see Methods, 33). Binding kinetics obtained from this method were compared with those from classical bilayer recordings.

### Kinetic rate constant of short peptides-OmpF channel interaction

Single-channel recordings using automated patch was employed to determine the binding kinetics of cationic peptides interacting with the OmpF channel. The applied transmembrane voltage played a critical role in modulating their interaction dynamics because of the positive charge of the peptides (30, 31).

At a transmembrane potential of -100 mV, transient current blockades were observed with two distinct durations: *τ*_*off*-0_ (short-lived events) and *τ*_*off-1*_ (long-lived events). The short-lived blockades (*τ*_*off*-0_), attributed to transient collisions of peptides with the channel openings, were voltage-independent and excluded from kinetic rate calculations (20, 26). In contrast, the long-lived events (*τ*_*off-1*_) were voltage-dependent and indicative of substantial peptide partitioning into the channel pore lumen.

Dwell-time histograms were fitted using exponential models: a two-exponential fit for penta-lysine (*τ*_*off*-0_ = 0.10 ± 0.20 ms and *τ*_*off-1*_ = 0.83 ± 0.04 ms) and tetra-arginine (*τ*_*off*-0_ = 0.28 ± 0.05 ms and *τ*_*off-1*_ = 0.57 ± 0.45 ms), and a single-exponential fit for tri-arginine (*τ*_*off-1*_ = 0.09 ± 0.02 ms). *τ*_*off*-0_ corresponded to low-amplitude blockades, while *τ*_*off-1*_ represented high-amplitude, longer-duration events.

Event frequency decreased at lower transmembrane potentials, resulting in reduced association rate constants (*k*_*on*_) (Fig. 6A, C). Notably, penta-lysine showed a pronounced increase in normalized event frequency with increasing voltage (Fig. 6D). The dwell time *τ*_*off*_ associated with peak 1 decreased at higher voltages, consistent with voltage-enhanced dissociation. This behavior supports a voltage-dependent binding interaction between the peptides and the OmpF pore. In the absence of such interactions, *τ*_*off*_ decreased uniformly with increasing voltage.

**Fig. 6.**
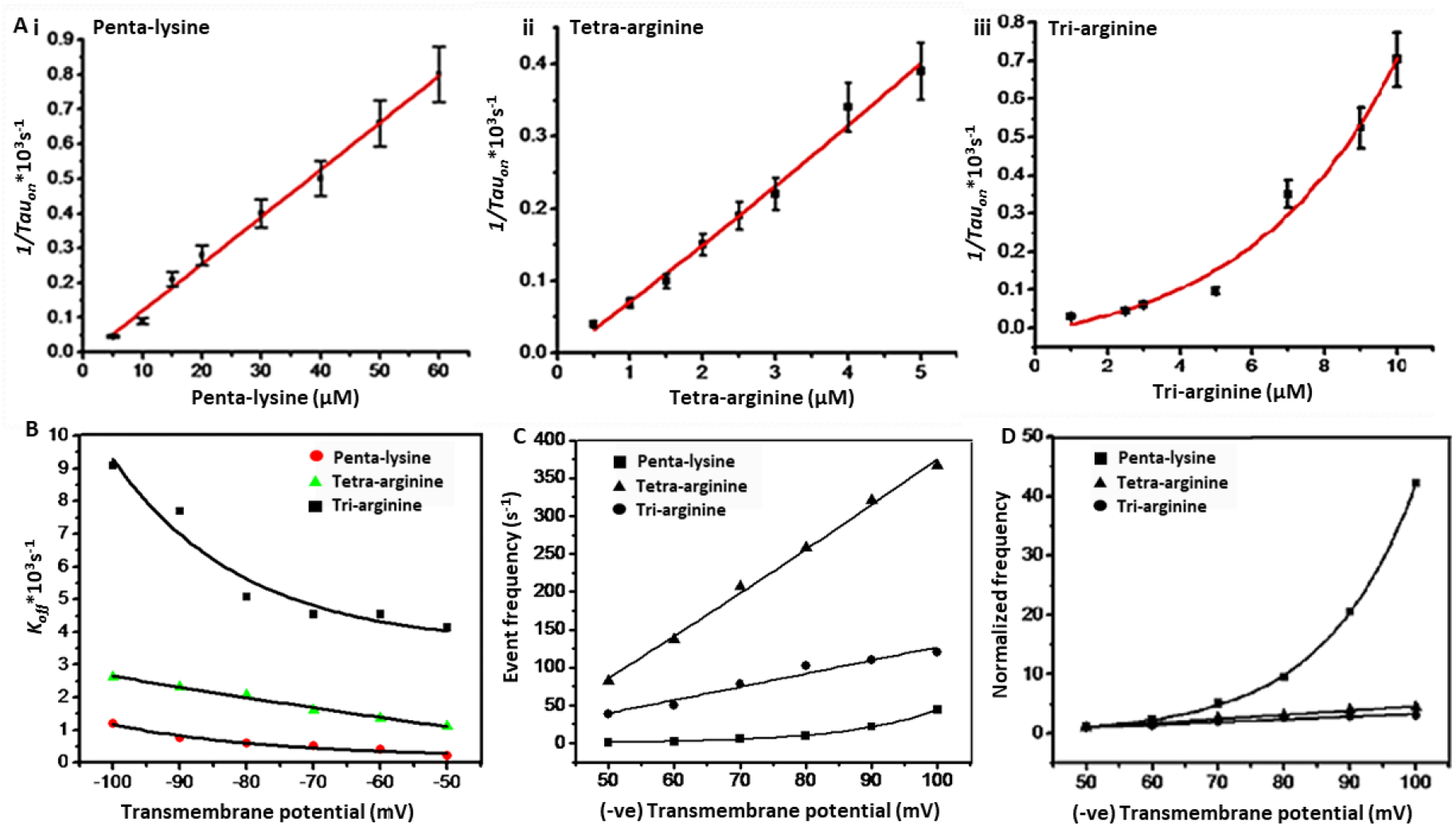
Concentration dependence of rate constant (*k*_*on*_) and voltage dependence of (*k*_*off*_) of peptides. (**A**) Association rate constant (*k*_*on*_) increased with increase in concentration of short polypeptides Penta-lysine (**i**), Tetra-arginine(**ii**) and Tri-arginine (**iii**) at applied potential of -100 mV. (**B**) Dissociation rate constant (*k*_*off*_) decreased with an applied increase in voltage. (**C**) Event frequency of the major peak 1 increased with an applied increase in voltage. (**D**) Normalized event frequency recorded at -50 mV. Experimental conditions: T = 25°C, 1 M KCl, 20 mM MES, pH = 6 and 5 µM concentration of peptides added to the cis side of the bilayer membrane.

Rate constants of the three peptides were derived from single-channel recordings at -100 mV in 1 M KCl (Table 1). The mean inter-event interval (*τ*_*on*_) was inversely proportional to peptide concentration, while *τ*_*off*_ remained concentration-independent (Fig. 6A), suggesting a simple bimolecular interaction model. The association rate constant (*k*_*on*_) was obtained from the slope of 1 / *τ*_*on*_ vs. peptide concentration [polypept], and the dissociation rate constant (*k*_*off*_) was calculated from the average of 1 / *τ*_*off*_ values. The equilibrium binding constant (*K*_*f*_ = *k*_*on*_ / *k*_*off*_) quantified peptide affinity for the channel. Additionally, the flux (*J*) of single peptide molecule/s crossing from the cis to trans side was estimated using *k*_*on*_ and the effective molar concentration gradient (*Δc*) (Table 1).

**Table 1:** Table showing kinetics of short cationic polypeptides binding with OmpF. Binding dependence is based on peptide molecular mass and peptide length, [pept]* = 0.215 M. The data were obtained by taking average of 2-3 individual measurements of each peptide at T = 25°C and in 1 M KCl, 20 mM MES, pH = 6.

| Concentration of peptides: 5 $\mu M$ | $K_{on} = 1/\tau_{on}[c]$ ($Ms$)$^{-1} \times 10^{-6}$ (Cis side, -100 mV) | $K_{on} = 1/\tau_{on}[c]$ ($Ms$)$^{-1} \times 10^{-6}$ (Trans side, 100 mV) | $K_{off} = 1/\tau$ ($s$)$^{-1} \times 10^{-3}$ | $K_f(M^{-1})$ | $J = K_{on} \Delta c/2(s)^{-1}$ (cis to trans) | $\Pi$ (Partition coefficient) = $K_f[pept]^*$ |
| --- | --- | --- | --- | --- | --- | --- |
| Penta-lysine | 9.2 | 10.45 | 0.86 | 10580.06 | 23; 26.12 | 2274.71 |
| Tetra-arginine | 79.40 | 100 | 1.75 | 4505.94 | 198.5; 250 | 968.77 |
| Tri-arginine | 19.70 | 49.26 | 10.10 | 1950.73 | 49.25; 123.15 | 419.40 |

In summary, cationic peptides interacted with the OmpF channel in a voltage-dependent manner, producing reversible current blockades. Penta-lysine and tetra-arginine peptides showed stronger binding and longer residence times than tri-arginine. Advanced bilayer techniques enabled detection of fast events which were missed by classical setups. Kinetic modeling revealed distinct association/dissociation behaviors, providing insight into selective peptide translocation through β-barrel pores.

## Discussion

The findings provide an insight into the interaction dynamics between cationic peptides and bacterial porin, specifically OmpF. Protamine exhibited voltage-dependent binding to OmpF, with significant blockage events occurring only under trans-negative potentials. The absence of translocation and presence of gating behavior suggest that protamine primarily interacts at the channel entrance, leading to transient or complete closure rather than passage through the pore (32). Blocking events induced by Ptm were not definite indicative of translocation through either porins, suggesting surface-level interaction rather than passage through the channel.

Furthermore, the mechanisms by which Ptm binds to OmpF or OmpD channels and exerts its antibacterial activity are not well established. Protamine’s secondary structure appears unstable, potentially unfolding into a rod-like conformation during interaction or attempted translocation (33). Protamine weaker interaction with OmpD channel, characterized by shorter blockage events, highlights channel-specific differences in peptide recognition. Mutational analysis of OmpF (D113N– E117Q) showed that negatively charged residues in the L3 loop were not essential for binding, suggesting alternative interaction sites.

Similar behavior was observed with longer peptides such as poly-arginine and poly-lysine, which induced mixed blockade events without evidence of translocation (34, 35). These results suggested that peptide-induced channel closure was a common mechanism contributing to antibacterial activity by disrupting membrane permeability.

Previous studies have shown that longer polypeptides interact similarly with α-hemolysin, a larger pore than OmpF (20, 36), with translocation properties influenced by peptide physiochemical characteristics (27). Our results with smaller cationic peptides such as penta-lysine, tetra-arginine and tri-arginine on OmpF revealed binding affinities and kinetic rate constants that depend not only on voltage and peptide length, but also on charge and molecular mass (37). Hydrophobicity and conformational flexibility (folded vs. unfolded states) may further influence interaction and transport across β-barrel pores (25, 31). The more hydrophobic the peptides were, the lower the rate constant of association was exhibited (38, 39).

Voltage-dependent changes in dissociation rate constants were observed, and the stability of the OmpF pore under extreme conditions enabled examination of peptide-pore interactions over extended periods (36). Although conformational transitions within the pore was not directly confirmed, uniform current blockades suggest that peptides adopt stable conformations during interaction.

The exchange of cationic α-helical arginine-based peptides between the bulk aqueous phase and the transmembrane β-barrel of OmpF at a single-molecule level was investigated. Structural and functional studies, including molecular dynamics simulations (40), may elucidate the precise binding mechanisms at atomic level and conformational changes involved in peptide-porin interactions.

## Conclusions

To conclude, the study demonstrates that the OmpF porin serves as a robust single-molecule platform for investigating cationic peptide interactions and translocation. High-resolution conductance measurements revealed voltage-, concentration-, and peptide length-dependent binding kinetics. The data provides insights into the translocation pathway and rate-limiting steps. These findings contribute to understanding peptide transport mechanisms and support the design of targeted antimicrobial peptides against bacterial pathogens.

## Contributions

S.P. designed the study, contributed to the experimental methodology, performed all the experiments, acquired, analyzed and interpreted the data. S.P. wrote the original draft and reviewed the manuscript.

## Acknowledgments

This work was supported by the EU-project MRTN-CT 2005-019335 (Translocation) to M.W. I thank Mahendran R Kozhjiampara for his feedback.

## Declaration of generative AI and AI-assisted technologies in the manuscript preparation process

During the preparation of this work no generative AI and AI-assisted technologies were used in order to rephrase text.

## Data availability

Supplementary information

