## Supplementary information for "Probing voltage dependence interaction of cationic peptides with bacterial porins at a single-molecule level"

**Sonal Prasad**

**Running title:** Cationic peptide–porin interactions

**Supplementary data contains:**

**-Supplementary Methods text**

**-Figures with captions (S1 to S4)**

**Chip-based automated patch clamp (Port-a-Patch)**

To achieve low-noise recordings and enable detection of fast binding events, a planar lipid bilayer with minimal capacitance was formed. Giant unilamellar vesicles (GUVs) were prepared using the electroformation method (1, 2) in an indium tin oxide (ITO)–coated glass chamber connected to the Vesicle Prep Pro system (Nanion Technologies). The ITO layers on the glass slides served as electrodes due to their electrical conductivity.

A lipid solution containing 5 mM DPhPC and 10% (mol/mol) cholesterol in chloroform was deposited onto the ITO-coated glass surface. Electroformation parameters (amplitude, frequency, duration, etc.) were controlled via the Vesicle Control software (Nanion Technologies).

Purified wild-type OmpF (1.5 mg/ml in 1% octyl-POE) was reconstituted into GUVs as previously described (3). The final porin concentration was adjusted to 10–20 nM, with detergent reduced to 0.001%. To remove octyl-POE, Bio-Beads were added, and the mixture was incubated overnight at 4°C. Bio-Beads were subsequently removed by centrifugation. The resulting proteo-GUVs were stored at 4°C which remained stable for up to one week.

For bilayer formation, 2–3 µl of the proteoliposome suspension was applied to a microstructured borosilicate glass chip (Port-a-Patch, Nanion Technologies) containing a ~1 µm aperture. Electrophysiological recordings were performed using the Port-a-Patch automated patch-clamp system. Measurements were conducted in 1 M KCl, 20 mM MES, pH 6.0.

Single-channel currents were recorded using an Axopatch 200B amplifier (Axon Instruments) connected to Ag/AgCl electrodes. Signals were filtered with a 4-pole low-pass Bessel filter at 10 kHz and digitized at 50 kHz using a Digidata 1440A digitizer. Peptides were added to the cis side of the bilayer, and ion current blockages were monitored to detect binding events (4).

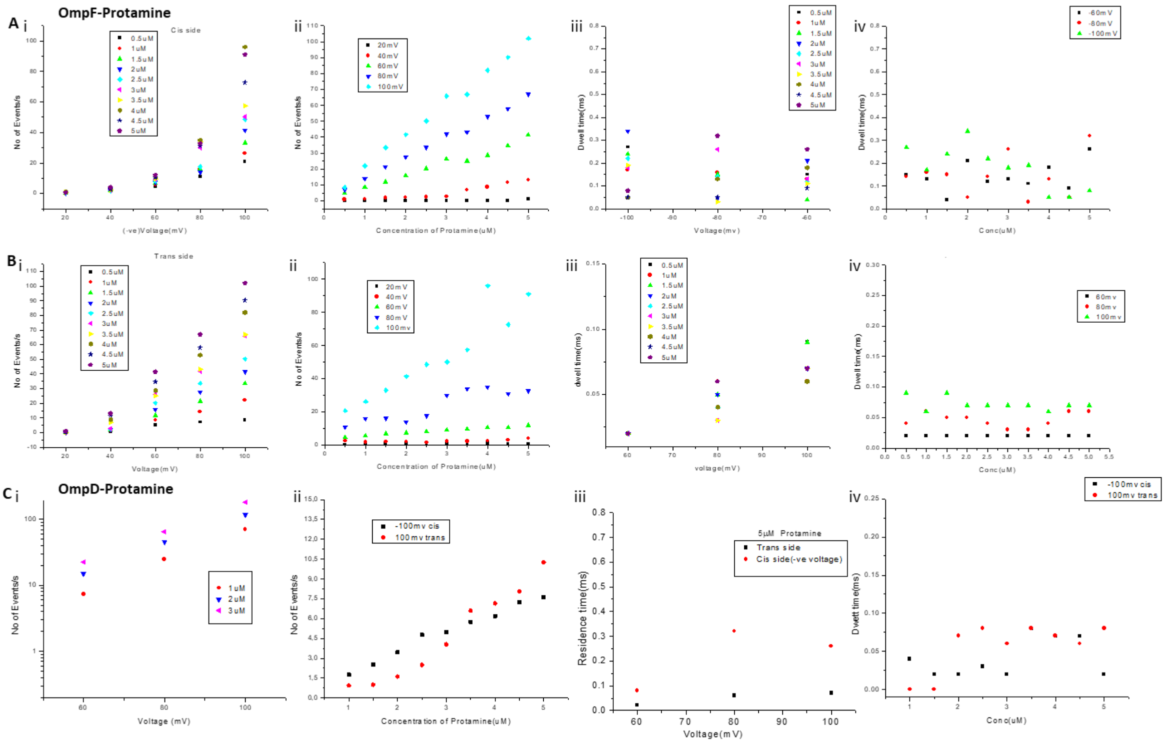


**Fig. S1. Concentration dependence of event frequency and voltage dependence of dwell time of Protamine through OmpF and OmpD porins**. (**A, B**) Event frequency (**i, ii**) and residence time (**iii, iv**) increased with both peptide concentration and applied voltage for OmpF channel after Protamine added to the cis and trans side. (**C**) Event frequency (**i, ii**) and residence time (**iii, iv**) increased with both peptide concentration and applied voltage for OmpD channel after Protamine added to the cis and trans side. Experimental conditions: T = 25°C, 1 M KCl, 20 mM MES, pH = 6.

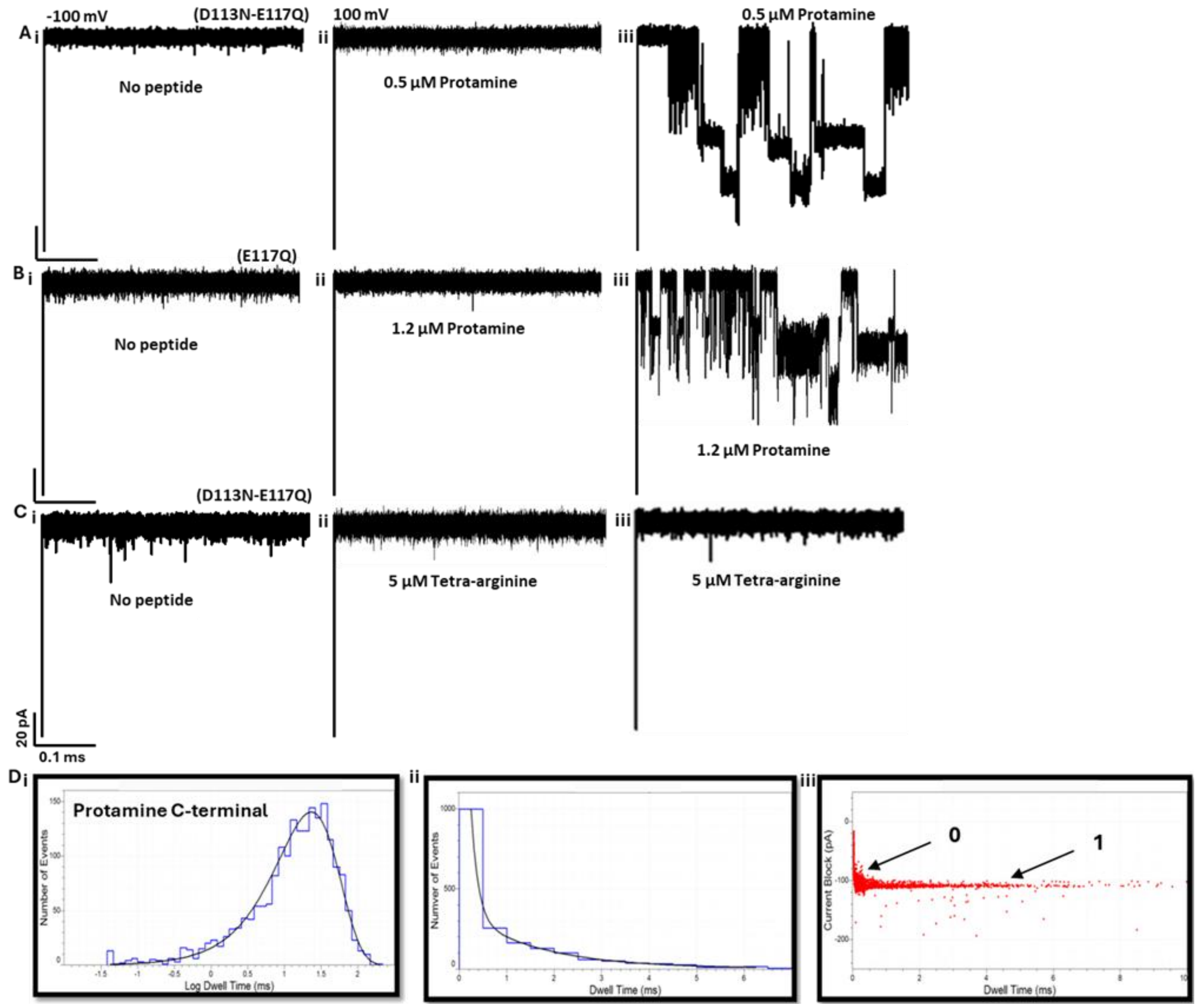


**Fig. S2. Typical ion current recordings through OmpF mutant porin with Protamine added to the cis side.** (**A, B, C**) In the absence of peptides at a transmembrane potential of -100 mV through single trimeric OmpF mutant channels reconstituted into planar lipid membranes (**i**). In the presence of peptides showing no interaction at a reversal transmembrane potential of 100 mV (**ii**). (**A, C**) Low concentration of protamine displaying inhibition of the double mutant and single mutant OmpF channels through two-step closure short blockage events at -100 mV (**iii**). (**B**) Tetra-arginine displayed no interaction with the double mutant OmpF channel suggesting specificity of Protamine interaction with OmpF porin (iii). (**D**) Semilogarithmic histogram of inter-event intervals of Protamine C-terminal end ($\tau_{on}$ = 29.87 ± 1.19ms), dwell-time histogram ($\tau_{off\text{-}0}$ = 0.10 ± 0.20 ms, $\tau_{off\text{-}1}$ = 0.83 ± 0.04 ms) and scatter plot of current block amplitude versus dwell time at an applied voltage of -100mV. Experimental conditions: T = 25°C, 1 M KCl, 20 mM MES, pH = 6.

### Liposome swelling assay

OmpF porin (50 ng/μl in 1% octyl-POE) was reconstituted into multilamellar liposomes following the procedure of Nikaido and Rosenberg (5). Liposomes were prepared from Escherichia coli total lipid extract (phosphatidylethanolamine, phosphatidylglycerol, and cardiolipin; 2 mg/ml) in the presence of 15% (w/v) dextran (molecular weight 40,000) to ensure vesicle entrapment. Size distributions were determined using a Nano-ZS ZEN3600 Zetasizer (Malvern Instruments). Control liposomes were generated in parallel without addition of porin.

Osmolarity of all test solutions was measured using an Osmomat 30 osmometer (Gonotec). For swelling assays, 30 µl of liposome or proteoliposome suspension was diluted into 630 µL of isotonic test solution prepared in 5 mM Tris-phosphate buffer, pH 7.4, within a 1 ml cuvette. The solution was gently stirred, and absorbance at 500 nm was recorded in kinetic mode using a Cary-Varian UV–Vis spectrophotometer, as described by Pagès et al, 2008. Swelling rates were calculated from the initial slope of the absorbance trace using $\Phi = (1/A_i)*dA/dt$ and averaged across at least two independent experiments, as previously reported (6).

#### Dynamic Light Scattering

In the below measurements, unilamellar liposomes using 1 mg/ ml of POPC were prepared using buffer solution 5mM KCl, pH-6.3 in water. In brief, measurements using 1-15 times dilutions were conducted at 20 °C in disposable cuvettes on a Malvern HTTPS Particle Sizer in backscattering configuration with a fixed scattering angle of 173°. Dynamic light scattering revealed a homogeneous liposome preparation. The size distributions of the obtained liposomes were determined by using a Zetasizer Nano (Malvern Instruments) and found to have a major peak at ~100 nm in diameter. This finding was in line with the preparation procedure for unilamellar liposomes, even if not all liposomes may be unilamellar. To reach the equilibrium, passive diffusion allowed on average one Ptm molecule to enter and the kinetic was likely limited by entry through the porin. The detailed protocol was followed as described in (7).

#### Zeta Potential Measurements

The same liposome preparations used for DLS were analyzed for zeta potential, reflecting contributions from both surface charge and the Donnan potential. Aliquots (1 ml) were transferred to disposable capillary cells (Malvern Instruments), and measurements were performed on a Malvern Zetasizer Nano Z at an applied electric field strength of 10 mV/cm.

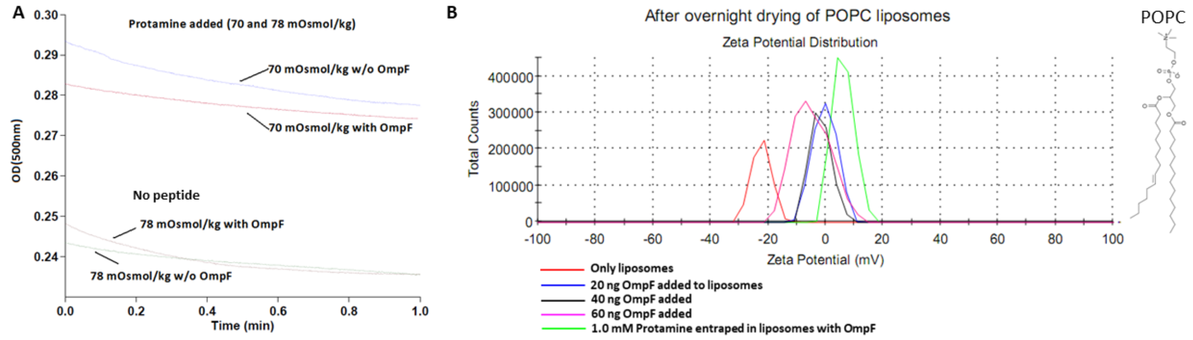


**Fig. S3: Size determination of the liposomes obtained by dynamic light scattering (DLS) and Zeta potential measurements**. (**A**) Liposomes (2.0 mM POPC lipid) reconstituted with OmpF porin showed increase in optical density (swelling rate) after addition of Protamine. Diffusion rate of protamine across liposome with OmpF was not high. The average size (diameter) of the liposomes was 97 nm, with a uniform size distribution that was desirable for kinetic investigations by tandem membrane assay later. (**B**) Increase in liposomes size integrated with OmpF size was observed after addition of Protamine. Small difference in Zeta potential in comparison of control liposomes with encapsulated liposomes with OmpF.

**Membrane Tandem Assay**

P-sulfonatocalix[4]arene (CX4), lucigenin (LCG), Triton X-100, octyl-polyoxyethylene (O-POE), protamine sulfate, and sodium phosphate were obtained from Sigma-Aldrich. POPC (1-palmitoyl-2-oleoyl-sn-glycero-3-phosphocholine; 25 mg/mL in chloroform) was purchased from Avanti Polar Lipids, and NAP-25 desalting columns from GE Healthcare. Purified OmpF (2 mg/ml in 1% O-POE) was used for proteoliposome preparation.

The CX4/LCG reporter assay was performed as previously optimized (8). Briefly, a lipid film was generated by depositing 100 µl of POPC stock solution (25 mg/ml) in a 5 ml round-bottom flask, followed by slow nitrogen drying and overnight vacuum desiccation. The film was hydrated with 1 ml of buffer containing 10 mM $NaH_2PO_4$, 700 µM CX4, and 500 µM LCG, and gently stirred on a rotary evaporator for ~20 min at room temperature. The resulting suspension underwent multiple times freeze-thaw cycles to generate unilamellar liposomes. Encapsulated CX4/LCG liposomes were separated from free dye using NAP-25 size-exclusion chromatography under identical buffer conditions (8).

For protamine and short-peptide uptake assays, 20 µl of CX4/LCG-loaded liposomes (25 µM lipid) in 10 mM sodium phosphate buffer (pH 7.0, 25 °C) were placed in a 2 ml cuvette, and time-resolved fluorescence was recorded. OmpF was added to a final concentration of 45 nM (2.8 µl of 2 mg/ml stock) to generate proteoliposomes in situ. At the end of each kinetic trace, Triton X-100 (50 µl of 1% solution) was added to lyse vesicles and determine maximal fluorescence. Fluorescence measurements were performed on a Varian Eclipse spectrofluorimeter using excitation at 367 nm and emission at 500 nm. Experimental details followed (7).

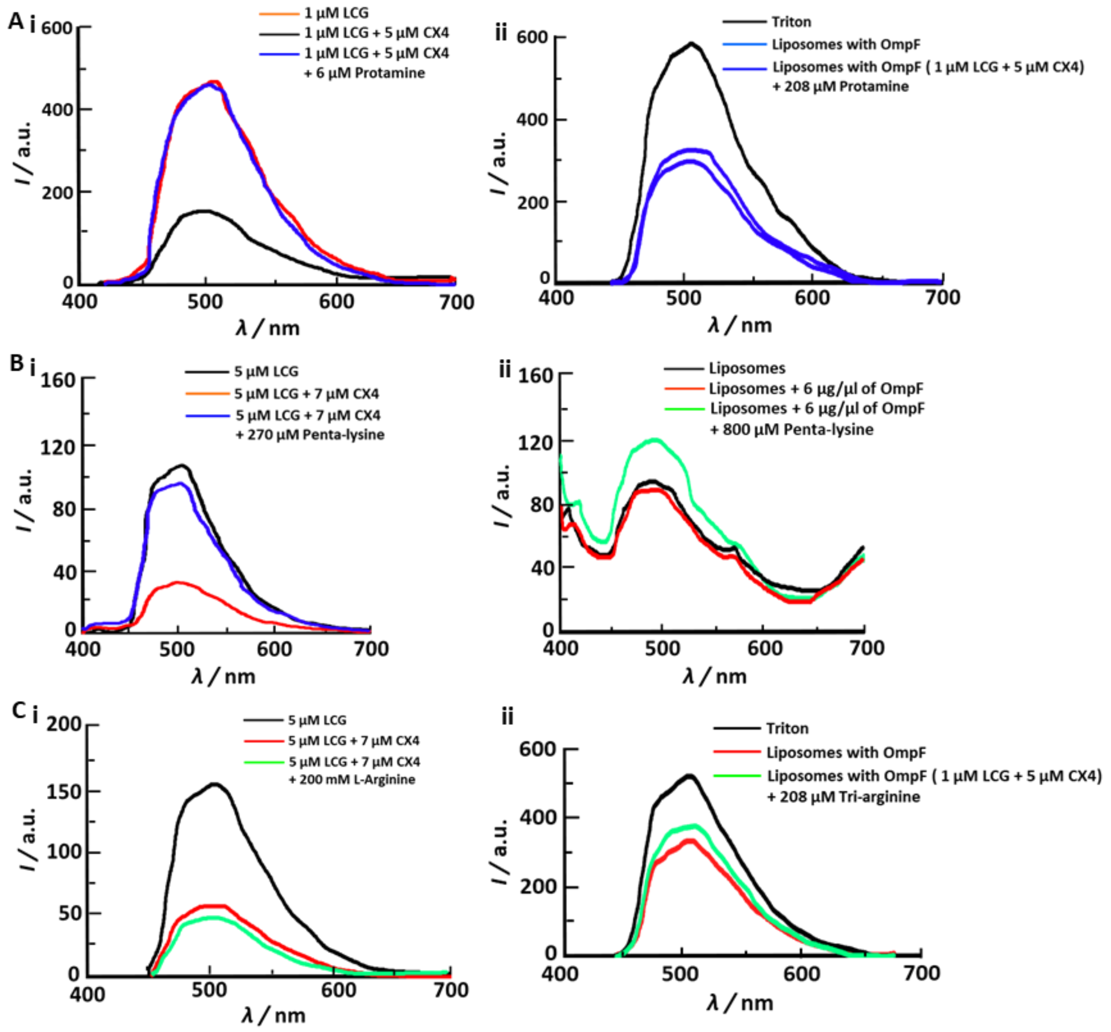


**Fig. S4: Substrate selective supramolecular tandem assay.** (**A, B, C**) Complete displacement of LCG dye from CX4 cavity by Protamine, Penta-lysine and Arginine (**i**). (A, B, C) Diffusion of the dye LCG and the host CX4 through the lipid membrane into the liposomes and further entrapment of host-guest and addition of OmpF and peptides. Protamine, Penta-lysine and Tri-arginine entered the liposomes via OmpF channels (**ii**).
